# Issues in the statistical detection of data fabrication and data errors in the scientific literature: simulation study and reanalysis of Carlisle, 2017

**DOI:** 10.1101/179135

**Authors:** Scott W. Piraino

## Abstract

**Background:** The detection of fabrication or error within the scientific literature is an important and underappreciated problem. Retraction of scientific articles is rare, but retraction may also be conservative, leaving open the possiblity that many fabricated or erroneous findings remain in the literature as a result of lack of scrutiny. A recently statistical analysis of randomized controlled trials [1] has suggested that the reported statistics form these trials deviate substantially from expectation under truely random assignment, raising the possiblity of fraud or error. It has also been proposed that the method used could be implemented to prospectively screen research, for example by applying the method prior to publication.

**Methods and Findings:** To assess the properties of the method proposed in [1], I carry out both theoretical and empirical evaluations of the method. Simulations suggest that the method is sensitive to assumptions that could reasonably be violated in real randomized controlled trials. This suggests that deviation for expectation under this method can not be used to measure the extent of fraud or error within the literature, and raises questions about the utlity of the method for propsective screening. Empirically analysis of the results of the method on a large set of randomized trials suggests that important assumptions may plausibly be violated within this sample. Using retraction as a proxy for fraud or serious error, I show that the method faces serious challenges in terms of precision and sensitivity for the purposes of screening, and that the performance of the method as a screening tool may vary across journals and classes of retractions.

**Conclusions:** The results in [1] should not be interpreted as indicating large amount of fraud or error within the literature. The use of this method for screening of the literature should be undertaken with great caution, and should recognize critical challenges in interpreting the results of this method.

## Introduction

Meta-research, a scientific endeavor aimed at studying and improving the process of science itself, has gained increasing interests among scientists. This interest has partially been driven by theoretical [2,3] and empirical work [4,5] that suggests concerns about the validity of the published scientific literature. One area of the scientific process that is amenable to meta-research is the detection of data validity/data integrity issues within the literature. Methods such as statcheck [6] and granularity testing [7] and its variants [8] have been developed to identify possible data validity issues by checking summary statistics reported in published research for consistency. In some cases, it has been proposed that the method be applied in an automated manner at various stages of the scientific process, for instance, prior to publication [6,9].

One class of methods for the detection of data validity issues is based on detecting whether data or summary statistics are consistent with their expected statistical distribution [10–15]. Under this framework, large deviations from the expected distribution of reported data are interpreted as indication of possible data integrity issues. In several cases within the literature, this method has been used to flag publications that were later determined to be based on fabricated data [11,12]. One variation on this, developed by Carlisle [11,13], uses reported summary statistics on baseline variables from randomized clinical trials to score published trials in terms of statistical deviation from that expected if subjects were truely assigned at random to various experimental groups. Large deviations potentially suggest issues with the validity of the reported summary statistics.

If methods for the detection of data validity issues are to play an increasing role in the scientific process, it is critical that scientists have a good understanding of the appropriate interpretation of these kinds of procedures. Of particular concern is that scientists may interpret the fact that a study or numerical result is flagged by these methods as substantial evidence of some type of flaw even when the method can sometimes flag an analysis for other reasons [16,17]. Especially if such methods are used to systematically screen research, it will be essential for scientists to have a grasp on the limitations that these methods may face. In order to understand these limitations, it is useful to distinqiush between multiple types of numerical results that may be identified by these methods. In what follows, I make a distinction between two different threats to data validity: data fabrication and data errors. Data fabrication may be said to occur when authors of published research intentionally alter data that they report in a way that is not consistent with how the data was collected or by reporting ficticious results about data that was never acutally collected. Data errors may be said to occur when authors of published research unintentionally report data in a way that is not consistent with what was acutally observed, for example by unintentional typographical errors or accidental errors in numerical calculations. Often, methods aimed at detecting data validity issues can be expected to flag both data fabrication and data errors, without distinqiushing between the two. In principle, this fact does not preclude the use of these methods for screening scientific research, because both fabrication and honest errors should be detected and corrected. However, the fact that these methods can not distingiush between errors and fabrication presents important interpretational challenges, since parties involved in the process are likely to respond differently if they interpret a flag by one of these methods as evidence of fabrication vs evidence of error. As a result, it is critical to manage expectations about what these methods show and how they should be applied.

Potentially of more concern for the application of these methods is the possiblility that some of the numerical results flagged by these methods are not erroneous, or in other words, are false positives. This may happen when there are aspects of data generation or reporting which are entirely legitimate but which are not accounted for by the method used to detect data validity issues. For example, it has been suggested that statcheck, which focuses on p-values, may result in false positives in cases where p-values are corrected (e.g. for multiple comparisons) but this correction is not taken into account [16,18]. Numerical results that are flagged as a result of these types of benign issues are problematic from a screening perspective because they can result in a waste of resources if all flagged analysis are investigated for potential data validity issues, as well as bringing unfair suspicion upon honest scientists. Understanding the relative frequencies of these different categories: fabrication, honest errors, and false positives, among flagged results is essential for the proper interpretation of these methods.

Although these issues are generally applicable to methods aimed at identifying data validity issues, they are particularly timely in light of a recent analysis by Carlisle [1], which applied a data validity detection method to a large sample of randomized controlled trials. This analysis has already generated significant attention both within the scientific literature [9] as well as the in the press [19,20]. The importance of [1] can be seen as relating to two related issues:

First, deviation from the expected statistical distribution of results across many clinical trials has implications for the global rate of fabrication or errors within the literature. This interpretation is apparent in the coverage of [1] (see [9], speculating that the results of [1] possibly indicate a “tsunami” of previously unrecognized fabrication in the literature, or [19], addressing the possible freqency of fabrication in literature based on figures from [1]). Fabrication and errors may be difficult to detect and may persist un-noticed in the literature [21], leaving open the possiblity that these ocurrences are not as rare as scientists might hope or desire [17,22]. This fact, combined with the observation that automated methods for error detection sometimes flag large proportions of the literature compared to what might be expected is potentially alarming. Certainly it appears that some observers have considered this interpretation [9,19,20]. The possiblity that a large proportion of of the scientific literature contains errors or fabrication would raise important question for the scientific community, and the suspicion that this is the case likely underlies part of the recent interest in meta-research. As a result, understanding the implications of methods and results such as those presented by Carlisle [1] for the overall rate of data validity issues in the literature is highly relevant.

Second, the method utilized by [1] is already being used to screen papers submitted to Anaesthesia [9], the journal in which [1] was published. The appropriate interpretation of methods for the detection of data validity issues in terms of screening the published literature, either retrospective or particularly prospectively (e.g. as a condition of publication) is an essential questions that remains to be addressed in the meta-research literature. Additionally, the editors of Anaesthesia have decided [9] to contact the journals associated with trials flagged by the analysis in [1], suggesting the need for serious investigations into these papers on the basis of the analysis in [1]. This suggests that understanding of the appropriate interpretation of [1] is urgently needed. In this article, I analyze the method utilized by Carlisle [1] to give insight into how it should be interpreted and what conclusions can be drawn from about these two critical questions.

## Results

To facilate understanding of the theoretical and empirical results I present, I briefly review the method utilized by Carlisle (which I refer to as the CM) [1]. For a single randomized controlled trial, the CM first involves manually extracting summary statistics on baseline (pre-treatment) variables from all groups which are randomized. For each variable, a p-value is calculated which tests the null hypothesis that the population means of the variable are equal across the groups. If the groups were truely assigned at random, then the null hypothesis is expected to be true for all of the variables. To combine the tests for all variables, the CM as applied in [1] utilized several methods for combining p-values that test a common null hypothesis, but [1] focuses on Stouffer’s method [23], which transforms the p-values to z-scores and calculates their sum. Under the assumption that the p-values included are independent, this sum is then compared to its own null distribution to derive a global p-value. Below, I highlight several stages at which this process may go wrong, along with re-analyses of the data used in [1] showing that these issues plausibly effected the analysis.

### Calculation of variable-level p-values from summary statistics

The CM as implemented in [1] involves calcuating p-values for for the differences in means of individual baseline variables within each trial using summary statistics, and then aggregating the p-values across each trial. Issues in the calcuation of the p-value for each variable may impact the validity of the downstream analysis. In order to test the ability of the method used by Carlisle [1] to recalculate p-values from summary statistics, I simulate data from two identically distributed groups and apply two of the p-value calculation methods used by Carlisle, a Monte Carlo method and ANOVA. The null hypothesis is true in these simulations, so the distribution of p-values should be uniform. Deviations from uniformity could indicate problems with the recalculated p-values, and could explain the deviations from uniformity that Carlisle [1] observed. To assess the robustness of these methods to assumption violations, I include two potentially problematic issues as part of my simulations. First, I simulate data from log-normal distributions instead of normal, as assumed in Carlisle’s analysis [1,13]. Second, I include rounding of reported summary statistics.

Fig 1 presents the distribution of simulated p-values. P-values generated from data with an underlying log-normal distribution and rounding to 2 digits (Fig 1A-C) display a roughly uniform distribution. Following Carlisle [1], I consider the closest p-value to 0.5 from multiple methods. When the underlying distribution is log-normal with 2 digit rounding this p-value has a slight excess near the center of the distribution compared to uniform. When some of the p-values are subject to extreme rounding (Fig 1D-F) the distributions display large excesses of p-values compared to uniform either near 1 (for ANOVA (Fig 1D)) or near 0.5 (for Carlisle’s Monte Carlo method (Fig 1E) or the closest to 0.5 of ANOVA and Monte Carlo (Fig 1E)). The observed p-values (Fig 1G-I) from [1] display some of these properties, with some large spikes of p-values near certain values, as well as possibly some excess in the center of the distributions.

**Fig 1:**
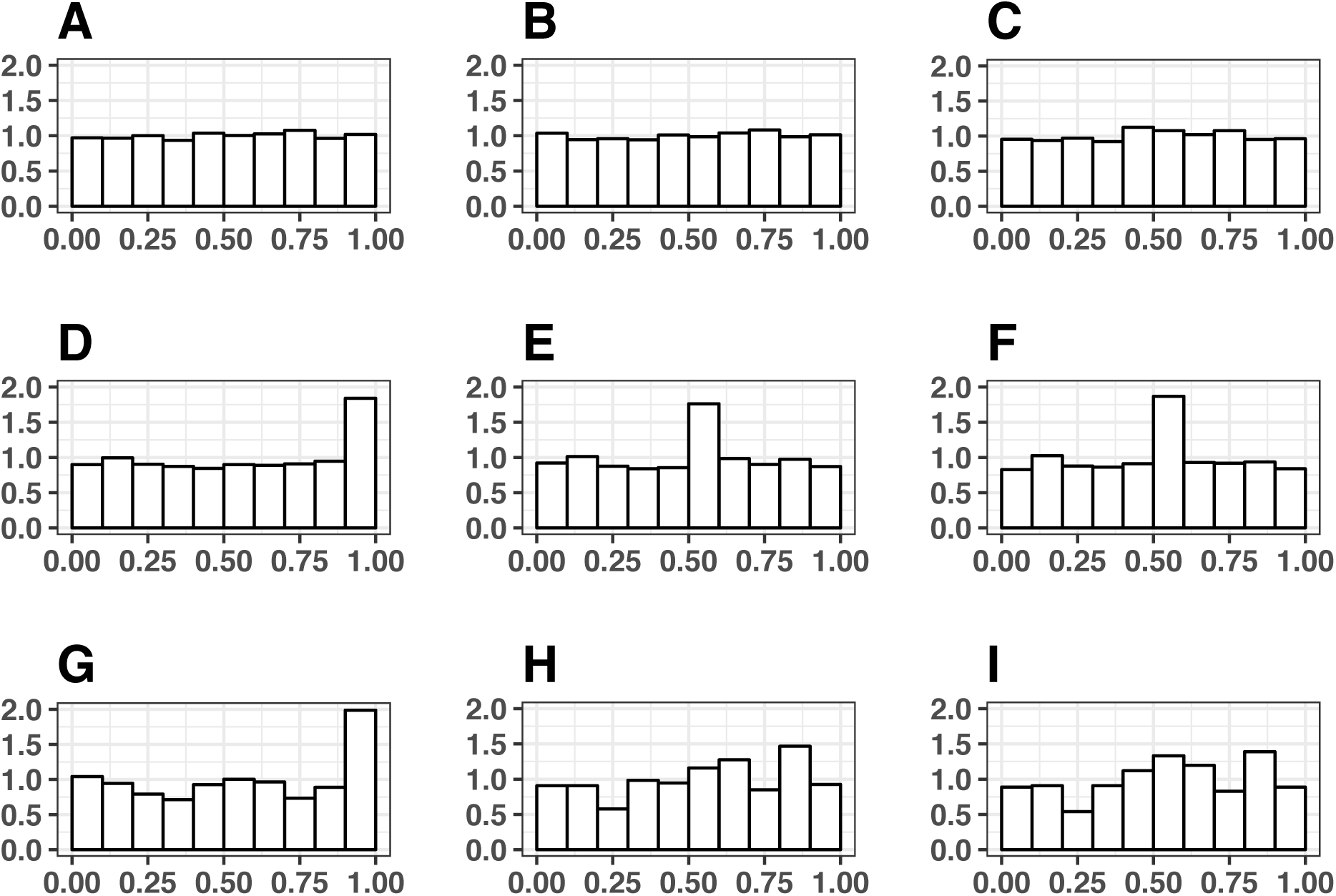
Distribution of simulated (A-F) and observed (G-I) p-values. The first row shows simulated p-values from summary statistics generated from log-normal distributions with moderate rounding for ANOVA (A) Carlisle’s Monte Carlo method (B) and the closed of those two to 0.5 (C). The second row shows simulated p-values from the model with summary statistics generated from a normal distribution where 90% of statistics have moderate rounding and 10% have extreme rounding for ANOVA (D) Monte Carlo (E) and the closest of the two to 0.5 (F). The third row shows observed p-values from the data collected by Carlisle from the Journal of the American Medical Association (JAMA) for ANOVA (G) Monte Carlo (H) and the closest of the two to 0.5 (I). FOr all rows, the first column shows ANOVA p-values, the second Monte Carlo p-values, and the third the closest of ANOVA and Monte Carlo to 0.5.

### Factors effecting trial-level p-values

Even if the variable-level p-values are validly calculated, issues may arise when multiple variable p-values are aggregated at the level of each trial. In Fig 2 I present simulations showing deviation from the expected null distribution of aggegated p-values (Fig 2A) under three conditions unrelated to data validity (Fig 2B-D). Fig 2B shows the distribution of trial p-values when the baseline variables that are aggregated are correlated with each other. The p-values show a pattern of excess p-values near 0 and 1, just as [1] observed. Fig 2C shows the distribution of p-values when there is imperfect randomization, resulting in residual confounding influence of the baseline variables. In this case, the p-values are right-skewed. Fig 2D shows the distribution of p-values when treatment assignment is randomized within strata that are associated with the baseline variables, resulting in left-skewed p-values. In all three cases, the p-value distributions have an excess of extreme p-values relative the the expected uniform distribution. If the CM is applied to a study for the purposes of screening, and one of these factors is application to the study, the CM may produce produce an extreme p-value for that study as a result of one of these factors rather than as a result of data validity issues. Likewise, for the global assessment of the prevalence of data validity issues in randomized control trials, deviations like those observed in [1] may be the result of a combination of these factors rather than a high prevalence of data validity issues.

**Fig 2:**
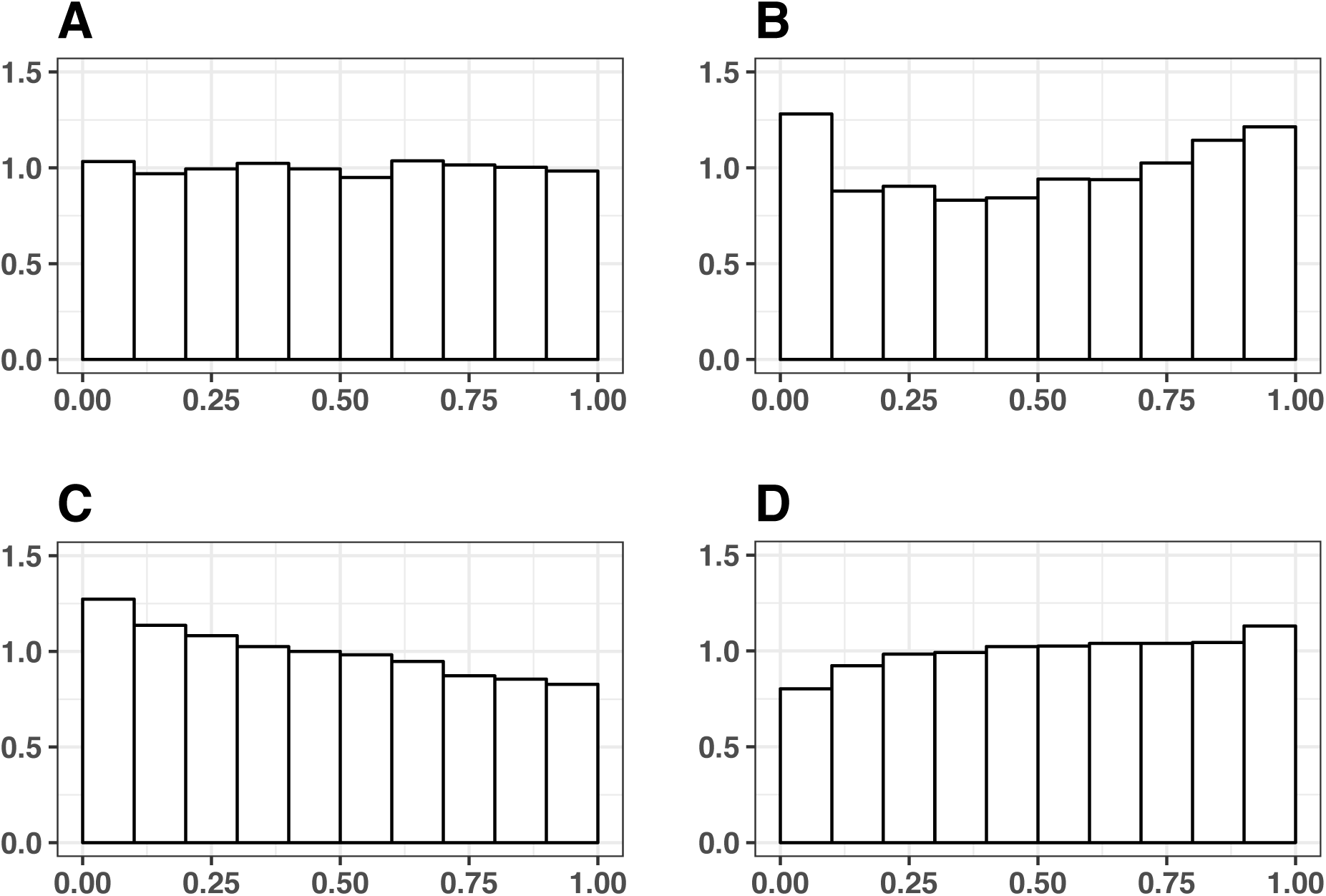
Histograms of p-values simulated from four different models. Null, with an expected uniform distribution (A) correlated baseline variables (B) imperfect randomization (C) and stratification (D).

### The Carlisle analysis is plausibly impacted by these issues

To determine if these issues plausibly played a role in the analysis conducted by Carlisle in [1], I reanalyzed the data from the supplement of [1]. Fig 3 compared theoretical distributions derived from simulations (Fig 3A and B, top row) with p-values from [1] (Fig 3C and D, bottom row). Fig 3A and B give the simulation distributions for null p-values and correlated baseline variable p-values, respectively. Fig 3C shows the distribution of p-values for all trials in [1], aggregated by Stouffer’s method. As Carlisle [1] notes, this distibution has an excess of p-values near 1 and 0 relative to the null (Fig 3A). However, this distribution is remarkably similar to the simulated distribution with correlated baseline variables (Fig 3B), suggesting that correlated variables could plausibly explain the deviations for uniformity.

**Fig 3:**
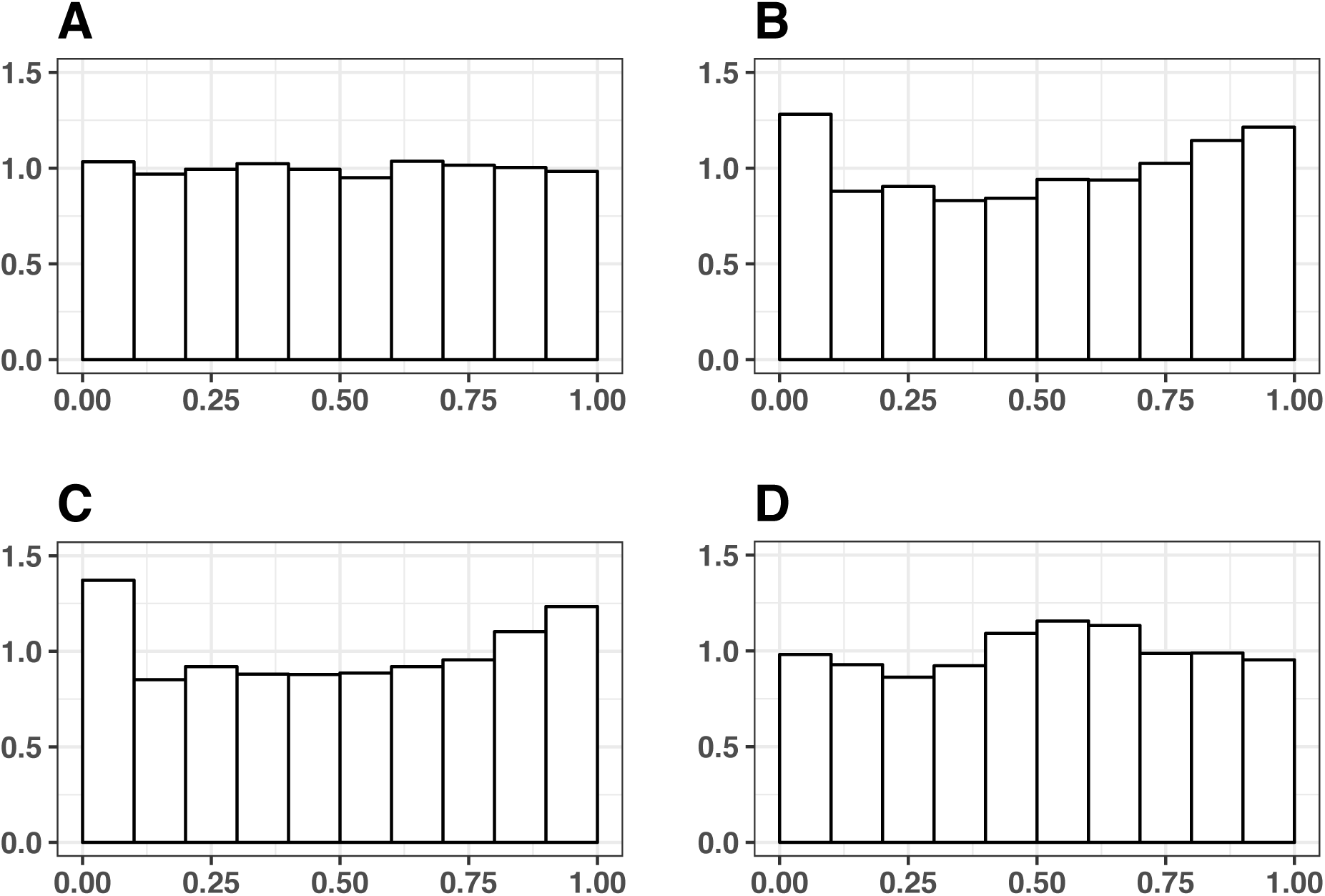
Distribtution of trial-level p-values for simulated null distribution (A), simulated p-values with correlated variables (B), observed trial-level p-values (C), and observed p-values for the first variable in each trial (D).

One objection to this is that the similarity between Fig 3B and C is sensitive to the simulation parameters. Indeed, I chose the parameters for the simulation intentionally to make the point that correlation can result in a similar p-value distribution. Other parameter settings can produce distributions which are less similar. In general, it is difficult to assess how realistic the simulation parameters are. For example, it could be argued that the correlation I used in this simulation (0.33) is higher than generally expected. On the other hand, this does not preclude correlation as an explanation for the results observed by Carlisle [1]. For instance, I assume all trials report 5 baseline variables. Trials that report more variables can have extreme deviations from uniform with lower correlations. Likewise, even if most correlations are lower, there may be some trials with extremely high correlation, or it could be that multiple factors (correlated variables, stratification, confounding) combine to form the observed distribution.

In general, it is difficult to definitively identify the cause of the deviation from uniformity using simulations alone. To overcome this, I examined the distribution of the first p-value for each trial (Fig 3D). Some causes of deviation from uniformity, such as fabrication or error, are expected to manifest on the level of individual variable-level p-values. Other causes, such as correlated variables, are expected to manifest when the p-values are aggregated. Comparison of the first variable p-values (Fig 3D) with the aggregated p-values (Fig 3C) can suggest what effects these different sets of explanations may have. The first variable p-values have a qualitatively different appearance compared to the aggregated p-values, lacking the excess of p-values near 0 and 1. This raises the possiblity that the excess extreme p-values are due to some issue with the aggregation process, rather than with the individual p-values themselves. The first variable p-values also display an excess of p-values in the center of the distribution relative to uniform. This may result from issues with the calculation of the individual p-values, as discussed above.

### Evalutation of the ability of the CM to identify retracted trials

The above analysis suggests that extreme trial-level p-values derived from the CM don not nessesarily indicate data validity errors. However, this does not nessearily preclude the usefulness of the CM for the detection of data fabrication and data errors. If the CM can identify known cases of fabrication or error in practice, then that empirical usefulness could form the basis for interpretation of the CM. Indeed, Carlisle analyzed retracted trials and showed that the CM p-values for retractions are more extreme compared to unretracted trials [1]. I extend this analysis by evaluation the distribtuions of the trial-level p-values from [1] across several retraction categories (Fig 4A and E).

**Fig 4:**
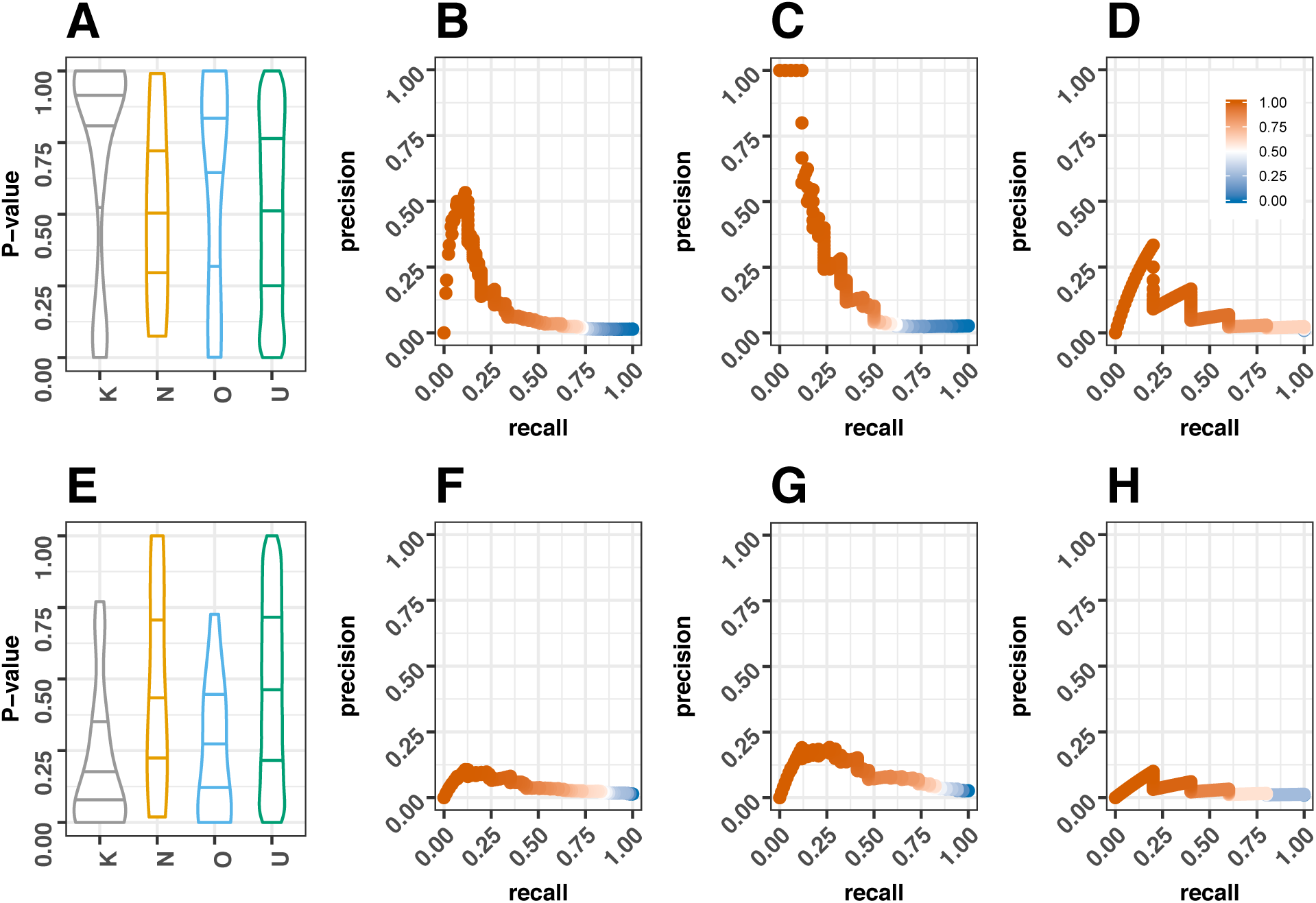
Assessment of the association between both one-sided (A-D, top row) and two-sided (E-H, bottom row) p-values with retraction status. Violin plots of one-sided (A) and two-sided (E) p-values for each of four trial catgories. Horizontal lines represent 25th, 50th, and 75th percentiles. The categories represent unretracted trials (“U”, green), trials retracted without indication of fabrication (“O”, blue), trials likely retracted for fabrication that were prominent examples known in the anesthesia community based on author names (“K”, grey), and trials likely retracted for fabrication that were not prominent examples known in the anesthesia community (“N”, orange). Panels B-D plot precision-recall curves for one-sided trial p-values for all trials (B), for trials in the journal Anesthesia and Analgesia (C), and for trials in the Journal of the American Medical Association (D). Panels E-G plot precision-recall curves for the inverse of the two-sided trial p-values (1 - p) for all trials (F), for trials in the journal Anesthesia and Analgesia (G), and for trials in the Journal of the American Medical Association (H). Color of the points in the precision recall curves indicate the threshold value used.

Using information contained in the supplemental materials of [1], I place each trial in one of four categories, based on it retraction status. I first divide the trials into those that have been retracted vs those that have not. I futher divide the retracted trials into three categories, starting by dividing them based on mention of fabrication in text descriptions of the retractions extracted by Carlisle. For those trials where fabrication is mentioned (and for which it is likely the reason for the retraction), I categorize the trials based on the presence of certain author names in the retraction descriptions. The CM has previously been used by Carlisle [11,13,15] to identify studies by several authors as potentially fraudulent. Several of these sets of studies are highlighted in the text of [1]. I seperately classify putatively fraud-based retractions based on the presence of these authors or other known authors of prominent anesthesia-related fraud cases to assess the possiblity that the association of trial-level p-values and retraction status differs between these groups.

Fig 4 shows the distribution of trial-level p-values across these four groups. Fig 4A shows the distribution of one-sided p-values, while fig 4E shows two-sided p-values. Consistent with the observations by Carlisle [1], trials by previously suspected anesthesiology-related authors (based on author name) (“K”, grey), and to a lesser extent trials retracted for putatively non-retraction related reasons both have abnormal p-value distributions, with the one-side p-values displaying an excess of p-values near 1 and a smaller excess near 0 (Fig 4A), and the two-sided p-values shifted toward 0 (Fig 4E). Both unretracted trials (“U”, green), and trials putatively retracted for fabrication that were not prominently known in the anesthesia community (“N”, orange) have p-value distributions much closer to uniform.

I also evaluate the ability of the trial p-values to identify retracted trials in terms of the precision (also called positive predictive value) and recall (also called sensitivity) of the p-values at various thresholds. I plot the results in precision-recall curves (Fig 4B-D and F-H), which displays precision-recall pairs when a p-value threshold is used to classify trials as retracted vs unretracted, for many different thresholds. The two-sided p-values (Fig 4 F-H, bottom row) show generally poor performance that is inferior to the one-sided p-values (Fig 4B-D, top row), so I focus further discussion on the one-sided p-values. As Loadsman and McCulloch [9] note in their commentary on [1], high recall (sensitivity) is not achieved without sacrificing precision. For all trials (Fig 4B, second column), precision is moderate, with the maximum slightly in excess of 0.5. Precision is also low at the highest p-values, suggesting that, while the retracted p-values are shifted towards 1, there are still unretracted trials that have high p-values as well. I also identified variability in the performance of the trial p-values across journals. For example, trials published in the jounrnal Anesthesia and Analgesia (Fig 4C) had precision near 1 for the highest p-values thresholds, while trials published in the Journal of the American Medical Association (Fig 4D) had poorer performance compared to the aggregate of all trials. The vast majority of trials are not retracted, which means that related to classification such as precision and recall can sometimes be based on small numbers of trials within particular categories. As a result, the results I present here should be considered with caution. Never the less, I beleive that these analyses raise important issues with regard to the applicability of the CM. The variablity in classification properties in different journals, along with the observed differences between fabrication previously identified by Carlisle vs new instances of fabrication, raises important questions about the generalizability of the CM, an issue which I dicuss more in depth below.

## Discussion

### Implications for global error rates

I first address the implications of the results I present here for the issue of global error rates. Readers of [1] may be concerned by the results presented there if they interpret the analysis to suggest that fraud or error are rampant in the literature, a possiblity that has already been aluded to by some observers [9,19,20]. The analysis that I have presented here indicates that the analysis by Carlisle [1] is not informative of the rate of data validity issues (either fabrication or error) within the literature. The pattern observed by Carlisle [1] in the global distribution of trial-level p-values can plausibly arise for benign reasons. When considering only a single p-values per trial, which avoids some of the problematic assumptions made in [1], the p-value distribution does not display the pattern that Carlisle identifies as potentially indicative of error, suggesting that this critique is not simply speculation.

### Implications for use of the CM for screening

The theoretical arguments I give also have implications for the use of the CM in screening. In particular, my theoretical results suggest that screening should not rely on probability statements based on the CM. For example, in [1], Carlisle sometimes thresholds the trial-level p-values (e.g. p < 1/10000). If the p-values produced by the CM were valid p-values, then it would be tempting to make statements like “p < 1/10000 would only happen once in 10,000 trials, if the trials were truely randomized”. My results suggest that these types of statements are not valid. For instance, using simulated p-values from correlated variables, 0.0024% of p-values are less than 1e-04, a 24 fold increase over the nominal rate.

If certain trials are particularly susceptible to these types of issues (e.g. a subset of studies with highly correlated variables, extreme rounding, strong stratification) this inflation could be exacerbated, without nessecarily being obvious to the user of the CM. Likewise, if multiple CM p-values are used together, as Carlisle [1] suggests could be done using multiple trials from the same author, the inflation of error rates compared to their nominal values could be further increased. For example, using correlated p-values, a single p-value of 0.01 has a p-value under correlation of 0.0291, a 2.91 fold inflation, while for two p-values of 0.01 combined by multplication, the inflation is 8.47 fold. This suggests that the p-values produced by the CM have a problem in terms of calibration. If users of the CM target a particular confidence level, in the presense of assumptions violations the p-values produced by the CM may not nessearily meet their nominal rates. In addition, extreme assumption violation may produce extreme p-values, so using conservative thresholds does not nessearily alleviate this problem.

In additional to problems with calibration, my analysis raises issues with the empirical performance of the CM in terms of its ability to classify known instances of error or misconduct. A global analysis, aggregating across types of retractions and across journals, indicates that when applying the CM there is a strong trade-off between precision and recall (sensitivity) as others [9] have speculated. This suggests that acceptable precision will result in low recall, suggesting that screening initiatives that utilize the CM may not result in significant proportions of errors being identified. If the CM generally identifies few errors, its benefits as a screening tool may be modest. In addition, even the optimal precision achieved by the CM in the full sample of trials is moderate. Retraction are rare, so a moderate precision does not imply that the CM is uninformative. However, there are important implications for the use of CM for screening. First, moderate precision warrants caution in the interpretation of results from the CM. Users should be aware that even at “conservative” thresholds, many of the flagged trials may not be erroneous or fraudulent. Second, parties that may consider using the CM for screening may consider false positives to be associated with increased costs of using the method, such as increased effort need to evaluate flagged trials or the potential of delaying publication of valid research over a false postive.

My analysis also reveals heterogeneity in the performance of the CM across categories of retractions and across journals. I discuss three possible explanations for this, all plausible. First, It may be that this heterogeneity arises from heterogeniety in the behavior of researchers who submit to different journals. If the processes by which error or fabrication occur tend to be different across the different journals and retraction categories, this could explain the observed heterogeneity.

Second, heterogeneity in precision-recall curves may arise due to issues in the detection of errors. Not all erroneous publications are retracted, and it may be that retractions are generally a conservative marker of error, such that there are many potentially erroneous trials within the literature that could go un-retracted. If this is, than the precision and recall rates calculated based on retraction could give a pessimistic picture of the CM. This is particually the case if several of the un-retracted trials that have extreme CM p-values are erroneous but undetected, which might be expected if the CM is an effective measure of error. Assuming this is the case, precision and recall using retraction as a metric may underestimate the values that would be obtained using the unseen labels of true error. Under this model, heterogeneity in precision and recall performance is really due to heterogeneity in the extent to which retraction detects error. This suggests the possiblity that the journals where the CM performs well are more indicative of the true performance of the CM, while the journals where it performs poorly simply underestimate performance because the trials with extreme p-values that remain un-retracted truely are erroneous, but simply have not been detected as such. Deeper looks at trials that produce extreme CM p-values are warranted to assess this possiblity.

Finally, it is possible the performance of the CM is overestimated in in the journals where it performs best. The journals where the CM performs well tend to be anesthesia journals that also contain retractions that may have been known to Carlisle during the development of the CM, and in some cases the journals contain retractions that occurred directly as a result of being identified by the CM. Retractions in this category show more extreme CM p-values compared to other fabrication-related retractions (Fig 4A and E). This suggests the possiblity that the CM may be “overfit” to these particular trials. If the CM was used to identify some retracte trials, it may be that erroroneous trials that have extreme CM p-values were more likely to be identified, while erroroneous trials that have less extreme CM p-values received less scrutiny within this sample, and therefore remain un-retracted. This may result in inflated recall values, due to the existence of un-retracted trials with moderate CM p-values.

This third possiblity may work synergistically with the first. For instance, if by chance the anesthesiology field happened to have several prominent examples of fabrication that display the property targeted in the CM, then it is possible that methods similar to the CM were more likely to emerge within this field. As a result, tests of these methods might be more likely to include anesthesia trials from this period, which happened to have an excess of trials displaying these properties, thus resulting in overestimation of the performance of these methods.

Taken together, this analysis suggests that caution is warranted if the CM is to be used for screening. Notably, some of the assumptions made my Carlisle are nessesitated by the fact that the analysis in [1] by nessessity was based on summary statistics. This significantly complicates the application for the CM. For example, addressing correlation among baseline variables would be very difficult using only summary statistics, but could be facilitated by analysis of the raw data. If journals choose to implement screening procedures prior to publication, authors of papers would be able to respond to the results of the CM by reanalyzing the raw data, thus potentially giving more definitive answers as to the validity of certain assumptions made by the CM. Critically, in order for this stategy to function well, authors, reviewers, and editors need to have a solid understanding of how various assumptions can impact the results of the CM. This paper can serve as a starting point for members of the scientific community who need to interpret results in this context. Likewise, when insitutions such as funders or journals consider whether or how the CM could play a role in decision making, the results presented here can give insight into the possible costs and benefits of various implementations.

## Methods

### P-value calculations

I recalculated p-values from summary statistics using the “anovaSummarized” command in the package “CarletonStats” [24], as well as a custom Monte-Carlo method used by Carlisle in [1]. To replicate the method used by Carlisle in my own simulations, I modified code provided by Carlisle in the comments at (http://steamtraen.blogspot.com/2017/06/exploring-john-carlisles-bombshell.html). I used the “metap” [25] package for combining p-values by Stouffer’s method [23] using the “sumz” function.

### Carlisle data

I obtained Data from the supplemental tables [1], and loaded the data into R using the “readxl” [26] package.

### Precision/recall analysis

I computed precision/recall curves using the package “PRROC” [27], with the CM p-value was the metric and a binary indictor of retraction (1 = retracted, 0 otherwise) as the target.

### Categorization of retractions

Table S1 of [1] contains notes by Carlisle with information about individual trials, including details of retractions. I Categorize the trials by detecting the presence of certain terms or word stems within these notes. I categorize a trial as having been retracted by the presence of “RETRACTED” within these notes, and un-retracted otherwise. I categorize a retraction as coming from a prominent anesthesia related author that is known for fabrication based on the presence of one of four names in the notes. I use the names “Sato”, “Boldt”, “Fujii” and “Reuben”. I catgorize a retracted trial that lacks one of these names as having been caused by fabrication based on the presence of “fabricat” in the notes. All other retracted trial I categorize as having ocurred for reasons other than fabrication. For one trial, manual review suggested that the note mentions “fabricat” but without definitively attributing the trial to fabrication. As a result, I label this trial as having occurred for reasons other than fabrication.

### Computational analysis

I conducted all analyses in R [28] version 3.3.2. I used the package “ggplot2” [29] for visualization.

### Reproducibilty and computational details

Code used to generate the analyses and figures included in this article are available at https://github.com/ScottWPiraino/carlisle_reanalysis.

